# Automatic grading of cervical biopsies by combining full and self-supervision

**DOI:** 10.1101/2022.01.14.476330

**Authors:** Mélanie Lubrano, Tristan Lazard, Guillaume Balezo, Yaëlle Bellahsen-Harrar, Cécile Badoual, Sylvain Berlemont, Thomas Walter

## Abstract

In computational pathology, predictive models from Whole Slide Images (WSI) mostly rely on Multiple Instance Learning (MIL), where the WSI are represented as a bag of tiles, each of which is encoded by a Neural Network (NN). Slide-level predictions are then achieved by building models on the agglomeration of these tile encodings. The tile encoding strategy thus plays a key role for such models. Current approaches include the use of encodings trained on unrelated data sources, full supervision or self-supervision. While self-supervised learning (SSL) exploits unlabeled data, it often requires large computational resources to train. On the other end of the spectrum, fully-supervised methods make use of valuable prior knowledge about the data but involve a costly amount of expert time. This paper proposes a framework to reconcile SSL and full supervision, showing that a combination of both provides efficient encodings, both in terms of performance and in terms of biological interpretability. On a recently organized challenge on grading Cervical Biopsies, we show that our mixed supervision scheme reaches high performance (weighted accuracy (WA): 0.945), outperforming both SSL (WA: 0.927) and transfer learning from ImageNet (WA: 0.877). We further shed light upon the internal representations that trigger classification results, providing a method to reveal relevant phenotypic patterns for grading cervical biopsies. We expect that the combination of full and self-supervision is an interesting strategy for many tasks in computational pathology and will be widely adopted by the field.

## 1 Introduction

Computational Pathology is concerned with the application of Artificial Intelligence (AI) to the automatic analysis of Whole Slide Images (WSI). Examples include cancer subtyping [5], prediction of gene mutations [26,15] or genetic signatures [15,8,16]. Predictive models operating on WSI as inputs, faces two main challenges: first, WSIs are extremely large and cannot be fed directly into traditional neural networks due to memory constrains. Second, expert annotations are laborious to attain, costly and prone to subjectivity. The most popular methods today rely on Multiple Instance Learning (MIL), which frames the problem as a bag classification task. The strategy is thus to split WSIs into small workable images (tiles), to encode the tiles by a NN and then to build a predictive model on the agglomeration of these tile encodings. Tile representations are crucial to the downstream WSI classification task. One common approach consists of initializing the feature extractor by pre-training on natural images, such as ImageNet. While such encodings are powerful and generic, they do not lie within the histopathological domain. Different strategies have thus been developed to reach more appropriate tile encodings using different levels of supervision.

A first strategy aims to learn tile features with full supervision [10,1]. For this, one or several experts manually review a large number of tiles and sort them into meaningful classes. While the model may thus benefit from medically relevant prior knowledge, the process is time consuming and costly. A second strategy consists of learning tile representations through self-supervision, leveraging the unannotated data. It has proven its efficacy [23,19] and even its superiority to the fully supervised scheme [7]. However, this approach has a non-negligible computational cost, as training necessitates around 1000 hours of computation on a standard GPU [7]. Moreover, it is not guaranteed that the obtained encodings are optimal for the prediction task we are trying to solve. While both techniques come have undeniable advantages, we hypothesized that combining them could allow us to benefit from the best of both worlds.

Motivated by these observations we propose a joint-optimization process mixing self, full and weak supervision (Figure 1). Our contributions are:

– We propose a method for mixed supervision that combines the power of generic encodings obtained from self-supervision with the biomedical meaningfulness obtained from full supervision.
– We measure the trade-off in performance between the number of annotations and the computational cost of training a self-supervised model, thus providing guidelines to train a clinically impactful classifier with a limited budget in expert and/or computational workload.
– With activation optimization (AM), we further show that the learned feature encodings extract biologically meaningful information, recognized by pathologists as true histopathological criteria for the grading of cervical biopsies. Of note, we demonstrate for the first time that AM can point to key tissue phenotypes. We thus provide a complementary method for network introspection in Computational Pathology.
– We used a cost-sensitive loss to boost the performance of grading by deep learning.

**Fig. 1.**
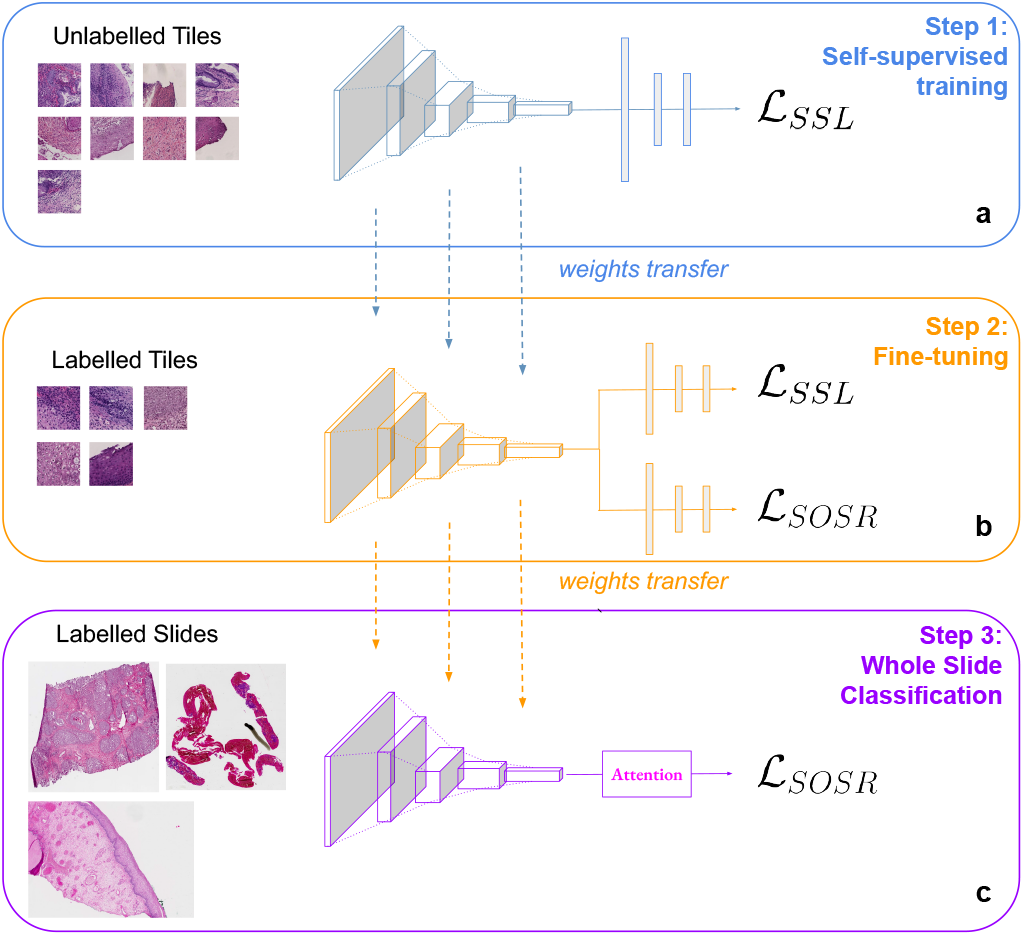
Mixed Supervision Process: **a)**A self-supervised model (SimCLR) is trained on unlabeled tiles extracted from the slides. Feature extractor and contrastive layer weights are transferred to the joint-optimization architecture **b)** Joint-optimization model is trained on the labeled tiles of the dataset. The feature extractor weights are transferred to the WS classification model. **c)** WSI classification model is trained on the 1015 whole slide images.

## 2 Related Work

Medical data is often limited. For this reason, one might want to take advantage of all the available data even if annotations might not be homogeneous and even though they might be difficult to exploit because multiple levels of supervision are available. AI applications have usually been dichotomized between supervised and unsupervised methods, spoiling the potential of combining several types of annotations. For this reason, mixing supervision for medical images analysis has gained interest in past years [13,18,17].

For instance, in [20] the author showed that combining global labels and local annotations by training in a multi-task setting, the capacities of the model to segment brain tumors on Magnetic Resonance Images were improved.

In [24], the author introduced a mixed supervision framework for metastasis detection building on the CLAM [19] architecture. CLAM is a variant of the popular attention based MIL [14] with 2 extensions: first, in order to make the method applicable in a multi-class setting, class-specific attention scores are learned and applied. Second, the last layer of the tile encoding network is trained to also predict the top and bottom attention scores, thus mimicking tile-level annotations. In [24], the authors highlight the limitations of this instance-classification approach and propose to leverage a low number of fully annotated slides to train the attention mechanism. In a second step, they propose to turn to a standard MIL training (using only slide-level annotations). Even with few annotated slides, this approach allows to boost classification performance and is thus the first demonstration of the benefits of mixed supervision in computational pathology. However, there are also some limitations. First, the method relies on exhaustive annotation of selected slides: for the annotated slides, all the key regions are annotated pixel-wise. Second, due to the CLAM architecture, the approach only fine-tunes a single dense layer downstream the pre-trained feature extractor. Third, the algorithm has been designed for an application case in which the slide and tile labels coincide (tumour presence). This however is not always the case: when predicting genetic signatures, grades or treatment responses, it is unclear how tile and slide level annotations relate to each other. In this article, we propose to overcome these limitations. We propose to combine self-supervised learning with supervision prior to training the MIL network. We thus start from more powerful encodings, that are not only capable of solving the pretext task of self-supervised learning, but also the medical classification task that comes with the annotated tiles. Consequently, this method does not require full-slide annotations, optimizes the full tile encoding network and does not come with any constraint regarding the relationship between tile and slide level annotations.

## 3 Dataset and Problem Setting

The Tissue Net Challenge [9] organized in 2020, the *Société Française de Pathologie (SFP)* and the *Health Data Hub* aimed at developing methods to automatically grade lesions of the uterine cervix in four classes according to their severity.

### Fully Supervised Dataset

5926 annotated Regions of Interest (ROIs) of fixed size 300×300 micrometers were provided. Each ROI had roughly the same size as a tile at 10x magnification and were labeled by the severity of the lesion it contained: normal (0) if tissue was normal, (1) low grade dysplasia or (2) high grade dysplasia if it presented precancerous lesions that could have malignant potential and (3) invasive squamous carcinoma.

### Weakly Supervised Dataset

The dataset was composed of 1015 WSIs acquired from 20 different centers in France at an average resolution of 0.234 +/− 0.0086 mpp (40X). The class of the WSI corresponded to the class of the most severe lesions it contained (grade from 0 to 3). Slide labels were balanced across the dataset.

### Misclassification Costs

Misclassification errors do not lead to equally serious consequences. Accordingly, a panel of pathologists established a cost matrix that assigns to each combination of true class and prediction (*i, j*) ∈ {0, 1, 2, 3}^2^ a severity score 0 ⩽ *C_i,j_* ⩽ 1 (Table 1). The metric used in the challenge to evaluate and rank the submissions was computed from the average of these misclassification costs. More precisely, if *P*(*S*) was the prediction of a slide *S* labelled *l*(*S*), the challenge metric *M_WA_* was:

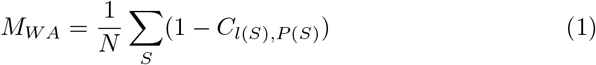

with *N* the number of samples.

**Table 1.** Weighted Accuracy Error Table - Error table to weight misclassification according to their gap with the ground truth

| Ground Truth | Benign (pred) | Low-grade (pred) | High-grade (pred) | Carcinoma (pred) |
| --- | --- | --- | --- | --- |
| Benign | 0.0 | 0.1 | 0.7 | 1.0 |
| Low-grade | 0.1 | 0.0 | 0.3 | 0.7 |
| High-grade | 0.7 | 0.3 | 0.0 | 0.3 |
| Carcinoma | 1.0 | 0.7 | 0.3 | 0.0 |

## 4 Proposed Architecture

### 4.1 Multiple Instance Learning and Attention

In Multiple Instance Learning, we are given sets of samples *B_k_* = {*x_i_*|*i* = 1 *.… N_k_*}, also called bags. The annotation *y_k_* we are given refers only to the bags and not the individual samples. In our case, the bags correspond to the slides, the samples to the tiles. We assume, that such tile-level labels exist in principle, but that we just do not have access to them. The strategy is to first map each tile *x_i_* to its encoding *z_i_*, which is then mapped to a scalar value *a_i_*, often referred to as attention score. The tile representations *z_i_* and attention scores *a_i_* are then agglomerated to build the slide representation *s_k_* which is then further processed by a neural network. The agglomeration can be based on tile selection [2,6], or on an attention mechanism [14], which is today the most widely used strategy.

### 4.2 Self-Supervised Learning

Self-supervised learning provides a framework to train neural networks without human supervision. The main goal of self-supervised learning is to learn to extract efficient features with inputs and labels derived from the data itself using a pretext task. Many self-supervised approaches are based on contrastive learning in the feature space. SimCLR, a simple framework relying on data augmentation was introduced in [3]. Powerful feature representations are learned by maximizing agreement between differently augmented views of the same data point via a contrastive loss 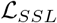 applied in the feature space. Details can be found in Supplementary Materials.

### 4.3 Cost-Sensitive Training

Instead of the traditional cross-entropy loss we used a cost-aware classification loss, the Smooth-One-Sided Regression Loss 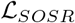. First introduced to train SVMs in [25], this objective function was smoothed and adapted for backpropagation in deep networks in [4]. When using this loss, the network is trained to predict the class-specific risk rather than a posterior probability; the decision function chooses the class minimizing this risk. The SOSR loss is defined as follows:

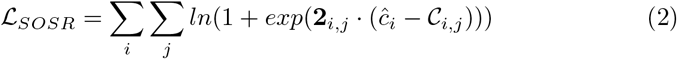

With **2**_*i,j*_ = −**1**_*i*≠*j*_ + **1**_*i*=*j*_, *ĉ*_*i*_ the *i*-th coordinate of the network output and *C* the error table.

### 4.4 Mixed Supervision

To be tractable, training of attention-MIL architectures requires freezing the feature extractor weights. While SSL allows the feature extractor to build meaningful representations [23,7], they are not specialized to the actual classification problems we try to solve. Several studies have shown that such SSL models benefit from specific fine-tuning to the downstream task [3]. We therefore added a training step to leverage the tile-level annotation and fine-tune the self-supervised model. However, as the final WSI classification task is not identical to the tile classification task, we suspect that fine-tuning solely on the tile classification task may over-specialize the feature extractor and thus sacrifice the generalizability of SSL (and for this reason ultimately also degrading the WSI classification performances). To avoid this, we developed a training process that optimizes the self-supervised and tile-classification objectives jointly. Two different heads, plugged before the final classification layer, are used to compute both loss functions 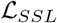 and 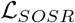 The final objective 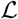 is then:

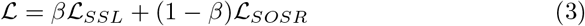

where *β* is a tuned hyperparameter. We found *β* = 0.3 (see Supplementary).

## 5 Understanding the Feature Extractor with Activation Maximization

To further understand the features learned by the different pre-training policies, we used Activation Maximization (AM) to visualize extracted features and provide an explicit illustration of the specificity learned. Methods to generate pseudo-images maximizing a feature activation have been introduced in [11]. This technique consists in synthesizing the images that will maximize one feature activation and can be summarized as follows [21]: if we consider a trained classifier with set of parameters *θ* that map an input image x ∈ ℝ^*h*×*w*×*c*^, (*h* and *w* are the height and width and *c* the number of channels) to a probability distribution over the classes, we can formulate the following optimization problem:

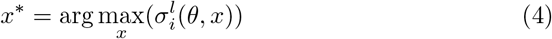

where 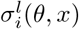 is the activation of the neuron *i* in a given layer *l* of the classifier. This formulation being a non-convex problem, local maximum can be found by gradient ascent, using the following update step:

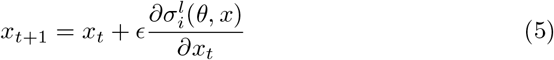

The optimization process starts with a randomly initialized image. After a few steps, it generates an image which can help to understand what information is being captured by the feature. As we try to visualize meaningful representations of the features, some regularization steps are applied to the random noise input (random crop and rotations to generate more stable visualization, details can be found in Supplementary Materials). To generate filter visualization within the HE space, we transformed the RGB random image to HE input by color deconvolution [22]. This preprocessing allowed to generate images with histology-like colors when converted back to the RGB space. To select the most meaningful features for each class, we trained a Lasso classifier without bias to classify the extracted feature vectors into the four classes of the dataset for the four pre-training policies. The feature vectors for each tile were first normalized and divided element-wise by the vector of features standard deviation across all the tiles. The L1 regularization factor *λ* was set to 0.01. Details about Lasso training can be found in Supplementary Materials. Contribution scores for each feature were therefore derived from the weights of the Lasso linear classifier: negative weights were removed and remaining positive weights were divided by their sum to obtain contribution scores ∈ [0, 1]. By filtering out the negative weights, the contribution score corresponds to the proportion of attribution among the features positively correlated to a class, and allows to select feature capturing semantic information related to the class, leaving out those containing information for other classes.

## 6 Experimental Setting

### 6.1 WSI Preprocessing

Preprocessing on a downsampled version of the WSIs was applied to select only tissue area and non-overlapping tiles of 224×224 pixels were extracted at a resolution of 1 mpp. (Details in Supplementary Materials)

### 6.2 Data Splits for Cross-Validation

To measure the performances of our models we performed 3-fold cross-validation for all our training settings. Because the annotated tiles used in our joint-optimization step were directly extracted from the slides themselves, we carefully split the tiles such that tiles in different folds were guaranteed to originate from different slides. The split divided the slides and tiles into a training set (70%), a validation set (10%) and a test set (20%). All subsequent performance results are then reported as the average and standard deviation of the performance results on each of these 3 test folds.

### 6.3 Feature Extractor Pre-Training

The feature extractor is initialized with pre-trained weights obtained with three distinct supervision policies: fully supervised, self-supervised or a mix of supervision. These three policies rely on the fine-tuning of a DenseNet121 [12], pre-trained on ImageNet. The fully-supervised archicture is fine-tuned solely on the tile classification task. The SSL architecture is derived from SimCLR framework and is trained on an unlabeled dataset of 1 million tiles extracted from the slides. Finally for the mixed-supervised architecture, a supervised branch is added to the previous SSL network and trained using the mixed objective function (see Fig. 1 and Eq. 3) on the fully supervised dataset. Technical details of these three training settings are available in the supplementary material.

### 6.4 Whole Slide Classification

After tiling the slides, the frozen feature extractor (DenseNet121) was applied to extract meaningful representations from the tiles. This feature extractor was initialized sequentially with the pre-trained weights mentioned above and generated as many sets of features. These bags of features were then used to train the Attention-MIL model with SOSR loss applied slide-wise. (Supplementary Materials).

### 6.5 Feature Visualization

To select the most relevant features, we trained an unbiased linear model on the feature vectors extracted from the annotated tiles. The feature vectors were standardised. The weights of the linear model were used to determine which features were the most impactful for each class. Feature visualizations were generated for the selected features and for each set of pre-trained weights (best training from cross-validation). We extracted the tiles expressing the most of these features by selecting the feature vectors with the higher activation for the concerned feature (Supplementary Materials).

## 7 Results

### 7.1 Self-Supervised Fine-Tuning

We saved the checkpoints of the self-supervised feature extraction model at each epoch of training, allowing us to investigate the amount of training needed to reach good WSI classification performances. We computed the embeddings of the whole dataset with each of the checkpoints and trained a WSI classifier from them. Figure 2 reports the performances of WSI classification models for each of these checkpoints. SSL training led to a higher Weighted Accuracy than using ImageNet weights after 3 epochs and resulted in a gain of +4.8% after 100 epochs. Interestingly, as little as 6 epochs of training are enough to gain 4% of Weighted Accuracy: a significant boost in performance is possible with 50 GPU-hours of training. We then observe a small increase in performance until the 100th epoch.

**Fig. 2.**
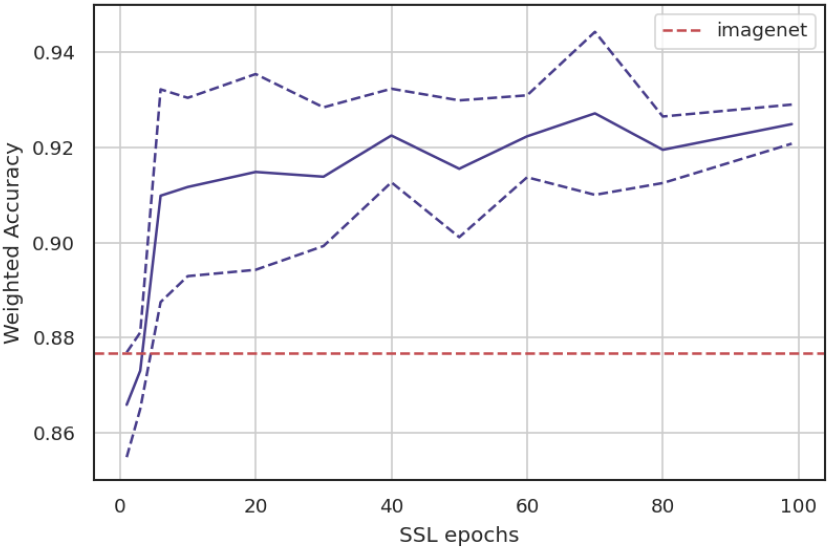
Weighted Accuracy (WA) for SSL and ImageNet pretraining. WA (obtained by 3-fold cross validation) as a function of the number of epochs for SSL training. Solid line: average WA, lines above and below: *±* standard deviation, respectively. Horizontal line: WA obtained by pretraining on ImageNet.

### 7.2 Pre-Training Policy Comparison

To compare the weights obtained with the various supervision levels, we ran a 3-fold cross-validation on the WS classification task and summarized the results in table 2. The results indicate that the SSL pre-training substantially improves the WSI classification performance. In contrast, we see that initializing the feature extractor with fully-supervised weights gives an equivalent or poorer performance than any other initialization. SSL pre-training allows us to extract rich features that are generic, yet still relevant to the dataset (unlike ImageNet). On the other hand, fully supervised features are probably too specific and seem not to represent the full diversity of the image data. The joint-optimization process manages to balance out generic and specialized features without neutralizing them: mixing the supervision levels brings significant improvements (+2%) to the performance, leading to a Weighted Accuracy of 0.945.

**Table 2.** Comparison of pre-training policies: performance depending on the pre-training regime (first column) and the classification loss (second column).

|  | Downstream Loss | Accuracy | Weighted Accuracy |
| --- | --- | --- | --- |
| ImageNet | Cross entropy | 0.758 +/- 0.034 | 0.865 +/- 0.023 |
| ImageNet | $\mathcal{L}_{SOSR}$ | 0.787 +/- 0.032 | 0.877 +/- 0.029 |
| Supervised | $\mathcal{L}_{SOSR}$ | 0.772 +/- 0.055 | 0.874 +/- 0.027 |
| SSL | $\mathcal{L}_{SOSR}$ | 0.803 +/- 0.016 | 0.925 +/- 0.006 |
| **mixed** | $\mathcal{L}_{SOSR}$ | **0.845 +/- 0.028** | **0.945 +/- 0.005** |

Next, we compared the cost-sensitive loss (Eq. 2) with the cross-entropy loss. Our results show that with ImageNet weights the SOSR loss improves the WA by 1% and the accuracy by 3%.

In conclusion, the combination of the SSL pre-trained model, its fully supervised fine-tuning, and the cost-sensitive loss leads to a notable improvement of 8 WA points over the baseline MIL model with ImageNet pretraining.

### 7.3 Number of Annotations vs Number of Epochs

We have seen that the combination of SSL and supervised pre-training lead to improved WSI classification. To further investigate the relationship between these two supervision regimes, we trained models with only some of the fully supervised annotations (15, 65, 100%) on top of intermediate SSL checkpoints. Results are reported in table 3. It appears that without SSL pre-training (or with too few epochs of training), the supervised fine-tuning does not bring additional improvement for WSI classification. This is in line with the work of Chen et al [3] that showed that an SSL model is up to 10x more label efficient than a supervised one. However, while fine-tuning the models by mixed supervision with too few annotations (15%) leads to a slight drop in WSI classification performances, we observe an improvement of 2 points of WA when using 100% of the tile annotations. Finally, we see a diminution of the standard deviations across splits for the different pre-training policies, showing better stability for longer SSL training and more annotations. We draw different conclusions from these observations:

– SSL is always preferable to only investing in annotations.
– The supervised fine-tuning needs enough annotations to bring an improvement to the WSI classification task. We can note however that even when considering the 100% annotation settings, the supervised dataset (approx. 5000 images) is still rather small in comparison to traditional image datasets.
– A full SSL training is mandatory to leverage this small amount of supervised data.

**Table 3.** Performance (WA) depending on SSL training time and number of annotations.

|  | 0 Annot. | ~1 Annot. / slide<br>(1015 tiles) | ~4 Annot. / slide<br>(3901 tiles) | ~6 Annot. / slide<br>(5926 tiles) |
| --- | --- | --- | --- | --- |
| **ImageNet (no SSL)** | 0,877 +/- 0.029 | 0.872 +/- 0.024 | 0.872 +/- 0.023 | 0,874 +/-0.027 |
| **SSL-epoch10** | 0,912 +/- 0.019 | 0,907 +/- 0.024 | 0,903 +/- 0.029 | 0,916 +/- 0.019 |
| **SSL-epoch50** | 0,915 +/- 0.014 | 0,913 +/- 0.024 | 0,916 +/- 0.014 | 0,914 +/- 0.022 |
| **SLL-epoch100** | 0,925 +/- 0.006 | 0,916 +/- 0.010 | 0,921 +/- 0.010 | **0,945 +/- 0.005** |

### 7.4 Feature Visualization

We generated the pseudo-images of the most important features for each class and each pre-training policy and extracted the related tiles. Figure 3 displays the most important features along with their most activating tiles for the class Normal (0). Although interpretation of such pseudo images must be treated carefully, we notice that the features obtained with SSL, supervised and mixed training are indubitably more specialized to histological data than those obtained with ImageNet. Some histological patterns, such as nuclei, squamous cells or basal layers are clearly identifiable in the generated images. The extracted tiles are strongly correlated with class-specific biomarkers. Feature **e** represents a normal squamous maturation, i.e. a layer of uniform and rounded basal cells, with slightly larger and bluer nuclei than mature cells. Features **c** and **d** highlight clouds of small regular and rounded nuclei (benign cytological signs). Feature **g** and **h** are characteristic of squamous cells (polygonal shapes, stratified organization lying on a straight basal layer). Interestingly, features extracted with the supervised method (**g**, **h**) manage to sketch a normal epithelium with high resemblance, the features are more precise. On the other hand, features extracted with SSL (**c**, **d**) highlight true benign criteria but do not entirely summarize a normal epithelium (no basal maturation). The mixed model displays both, suggesting that mixed supervision highlights pathologically relevant patterns to a larger extent than the other regimes [27].

**Fig. 3.**
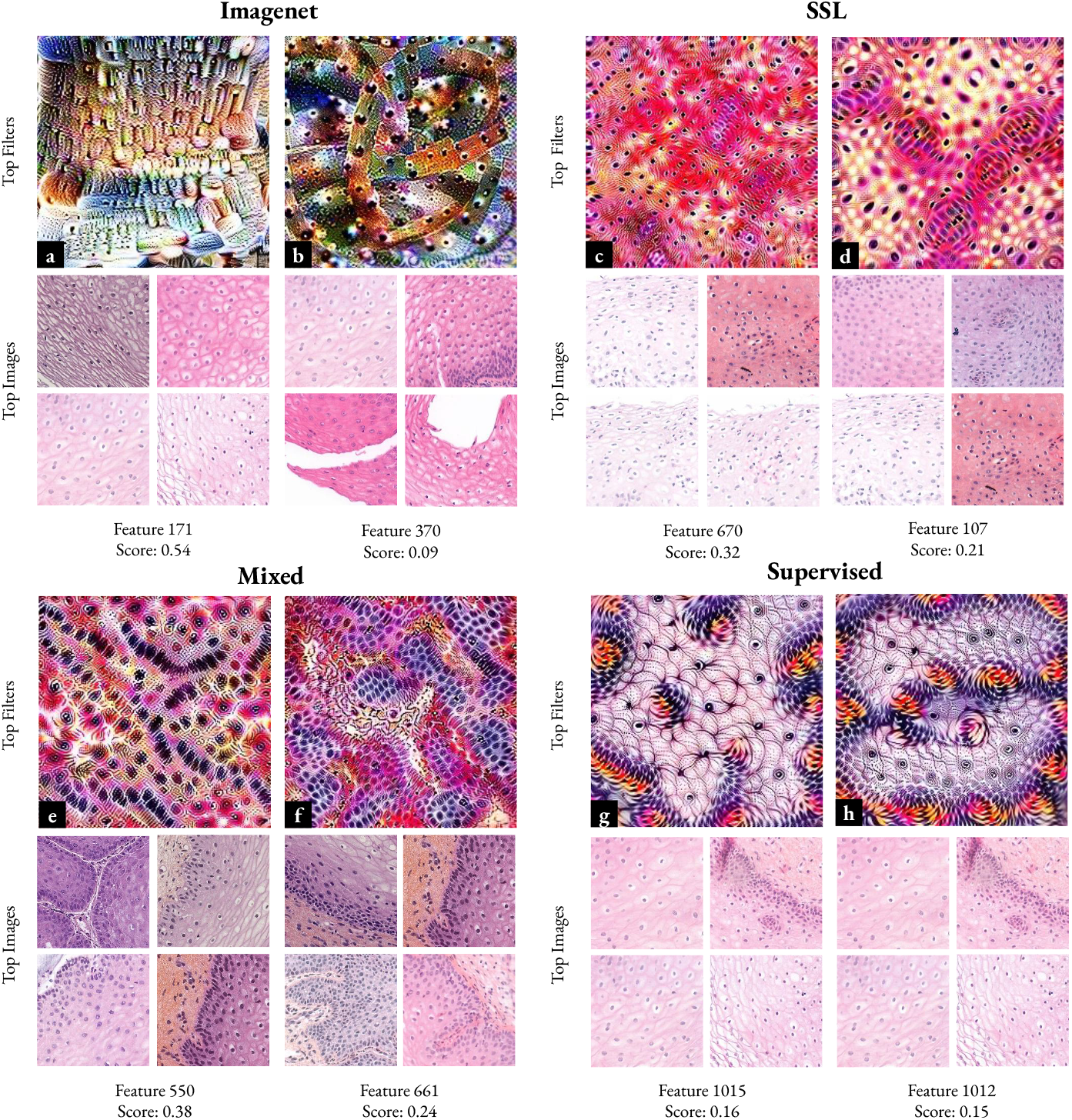
Feature Visualization. Top Features for class “Normal” (0) and tiles expressing the most the feature. Features obtained with SSL and Mixed Supervision are clearly related to histopathological patterns. Nuclei, squamous cells, basal layers and other histological morphologies are identifiable. Similar visualization for other classes are available in the Supplementary Materials.

In Figure 4 we can further identify class-related biomarkers for dysplasia and carcinoma grade. Tiles with visible koilocytes (cells with a white halo around the nucleus) have been extracted from the top features for Low Grade class. Koilocytes are symptomatic of infection by Human Papillomavirus and are a key element for this diagnosis (almost always responsible for precancerous lesions in the cervix, [27]). High Grade (2) generated image represents disorganised cells with a high nuclear-to-cytoplasmic ratio, marked variations in size and shape and loss of polarity. For the class Carcinoma (3), we observe irregular clusters of cohesive cells with very atypical nuclei, separated by a fibrous texture that can be identified as stroma reaction. All these criteria have been identified in [27] as key elements for diagnosis of dysplasia and invasive carcinoma.

**Fig. 4.**
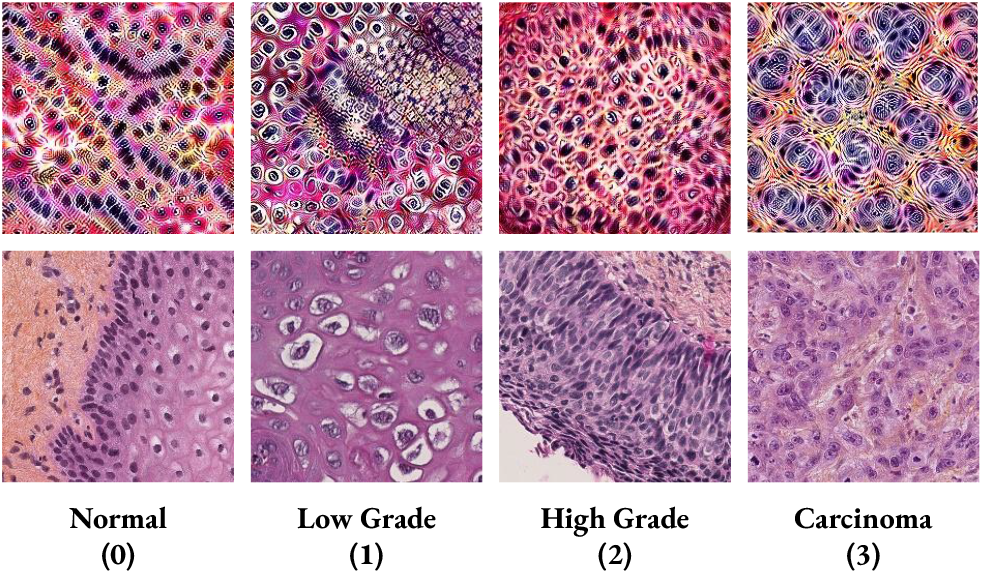
Feature comparison per class. The top row displays the top filter for the Mixed Supervised model for each class. The bottom row displays the tile expressing the feature the most. Extracted tiles correlate with class-specific biomarkers.

In Figure 5, we observe that features extracted from ImageNet and SSL models are diverse, in particular, features extracted from SSL reflect rich tissue phenotypes which correlates to their generic capacities of image representations. On the other hand, features extracted with supervised and mixed methods are more redundant. Visualizations for other classes are available in Supplementary Materials.

**Fig. 5.**
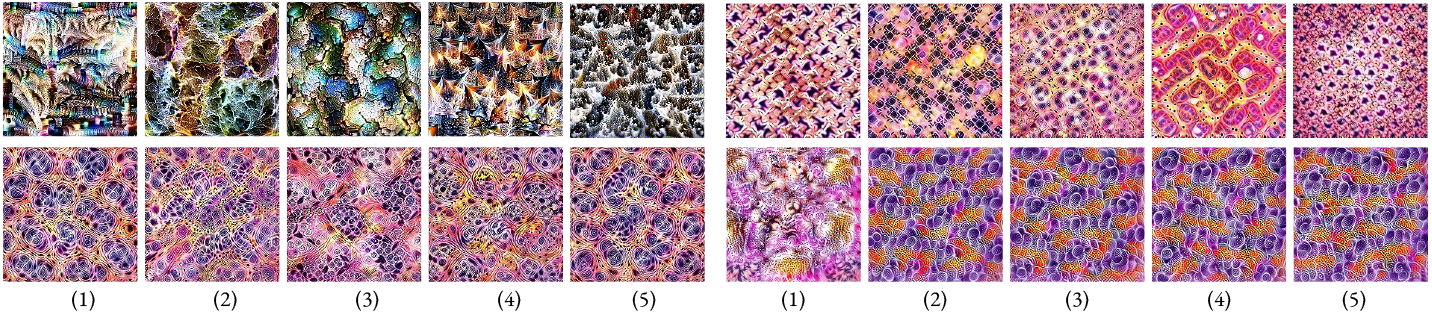
Feature Diversity for the class “Carcinoma” (3) (top 5 features) Top Left: ImageNet, Top Right: SSL. Bottom Left: Mixed, Bottom Right: Supervised.

## 8 Discussion

In pathology, expert annotations are usually hard to obtain. However, we are often in a situation where a small proportion of labeled annotation exists but not in sufficient quantities to support fully supervised techniques. Yet, even in small quantities, expert annotations carry meaningful information that one could use to enforce biological context to deep learning models and make sure that networks learn appropriate patterns. On the other hand, self-supervised methods have proven their efficacy to extract generic features in the histopathological domain and their usefulness for downstream supervision tasks, even in the absence of massive ground truth data. Methods capable of reconciling self-supervision with strong supervision can therefore be useful and open the door to better performance.

In this paper, we presented a way to inject the fine-grained tile level information by fine-tuning the feature extractor with a joint optimization process. This process thus allows to combine self-with full supervision during the encoder training. We applied our method to the TissueNet Challenge, a challenge for the automatic grading of cervix cancer, that provided annotations at the slide and tile level, thus representing an appropriate use case to validate our method of mixed supervision. We also propose in this study insights and guidelines for the training of a WSI classifier in the presence of tile annotations. First, we showed that SSL is always beneficial to our downstream WSI classification tasks. Fine-tuning pre-trained weights with SSL for only 50 hours brings a 4% improvement over WSI classification weighted accuracy, and near to 5% when fine-tuning for longer (100 epochs). Second, a small set of annotated tiles can bring benefit to the WSI classification task, up to 2% of weighted accuracy for a supervised dataset of around 5000 images. Such a set of tiles can be obtained easily by asking the pathologist to select a few ROIs that guided their decision while labeling the WSIs, which can be achieved without a strong time commitment. However this boost in performance can be reached only if the feature extractor is pre-trained with SSL, and for sufficiently long: SSL unlocks the supervised fine-tuning benefits.

To further understand the differences between the range of supervision used to extract tile features, we conducted qualitative analysis on features visualizations by activation maximization and observed that features obtained from SSL, supervised or mixed trainings were more relevant for histological tasks and that class-discriminative patterns were indeed identified by the model.

The scope of this study contains by design three limitations. First, SSL models were trained by fine-tuning already pre-trained weights on imagenet. This may explain the rapid convergence and boost in performance observed; however it may also underestimate this boost if the SSL models were trained from scratch. We did not compare SSL trained from scratch and fine-tuned SSL, and left it to future work. Second, all the conclusions reached are conditioned by the fact that we do not fine-tune the feature extractor network during the WSI classification training. Keeping these weights frozen, and even pre-computing the tile representations brings a large computational benefit (both in memory and speed of computations), but prevents the feature extractor from specializing during the WSI classification training. Third, the tendency observed in table 3 of better performances correlated with larger numbers of annotations is modest and would require more annotations to validate it. Application to different task such as rare diseases dataset with few data could show better benefit.

To conclude, we present a method that provides an interesting alternative to using full supervision, pre-training on unrelated data sets or self-supervision. We convincingly show that the learned feature representations are both leading to higher performance and providing intermediate features that are more adapted to the problem and point to relevant cell and tissue phenotypes. We expect that the mixed supervision will be adopted by the field and lead to better models.

## Acknowledgments

The authors thank Etienne Decencière for the thoughful discussions that help the project. ML was supported by a CIFRE PhD fellowship founded by KEEN EYE and ANRT (CIFRE 2019/1905). TL was supported by a Q-Life PhD fellowship (Q-life ANR-17-CONV-0005). This work was supported by the French government under management of ANR as part of the “Investissements d’avenir” program, reference ANR-19-P3IA-0001 (PRAIRIE 3IA Institute).

## Notes

### Competing Interest Statement

The authors have declared no competing interest.

### Summary of Updates

Change in affiliations (typo)

