## Supplementary Materials for "Automatic grading of cervical biopsies by combining full and self-supervision"


### 1 WSI preprocessing

To select only the tissue area of the WSIs, we enhanced the contrast of downsampled versions of the images, computed the standard deviation of each of the RGB channels, normalized the obtained standard deviation map and applied Otsu thresholding [1]. We then extracted non-overlapping tiles of 224x224 pixels at a resolution of 1 mpp.

### 2 Models implementations and training details

#### 2.1 Feature extractor pre-training

**Self-supervision** SimCLR architecture was used. It consisted of a feature extractor and a projection head. We chose DenseNet121 for the feature extractor followed by 3 dense layers for the projection head. We trained the SimCLR model with 1 million tiles (336x336 pixels) extracted from the slides of the dataset for 100 epochs on 8 GPUs. The training necessitated approximately 800 GPU hours. Weights were saved every epoch. We used a batch size of 864, a temperature of 0.1 and a learning rate of 10e-4. To boost the training we initialized the feature extractor with ImageNet weights. To make sure random noise from the projection head did not perturb imagenet features, we initialized the projection head by first training SimCLR for 3 epochs on 10 000 tiles while freezing the feature extractor. We then unfroze all the layers of the model and trained for 100 epochs on the 1 millions tiles with the contrastive loss (eq). The following data augmentation were used: crop and resize of the tiles to 224x224 pixels, rotation (90°), random flipping, custom stain augmentation and color jittering. Data augmentation parameters were carefully tuned to fit well with histology data (see details in table 3).

**Full Supervision** Image Classification network is summarized in table 1. The model was trained for 70 epochs with the SORS loss, using early stopping and keeping the best weights according to the validation performances. The tile classification model was trained on the 3 tiles splits generating 3 different sets of weights to be transferred ultimately to the whole slide classification model. Standard data augmentation was used (flips, rotation, translation, zoom, gaussian blur, color augmentation, see table 4) and the model was trained with a learning rate of 1e-4 and ADAM optimizer.

**Mixed Supervision** Mixed supervision architecture is summarized in table 1. The supervised branch was optimized with the SORS loss function. The parameter beta weighting the contribution of the 2 losses of the joint-optimization was optimized in a dedicated experiment and was set to 0.3. Similarly to the fully-supervised training, the mix-supervised model was trained on the 3 pre-defined tiles splits and generated 3 sets of weights accordingly. For each data split, the network was initialized with weights from SSL pre-training. So as not to disrupt feature extractor weights with initial random noise, the supervised branch was trained alone for two epochs. Then, all the layers were unfrozen and the network was trained for 200 epochs. Early stopping was applied and the best weights were kept.

| **Fully supervised** | **SSL** | **SSL + Supervision** | |
|---|---|---|---|
| **Feature Extractor**<br>DenseNet121<br>[1024] | **Feature Extractor**<br>DenseNet121<br>[1024] | **Feature Extractor**<br>DenseNet121<br>[1024] | |
| **Classification Head**<br>FC [128] + relu<br>Dropout<br>FC [64] + relu<br>FC [4]<br><br>$\mathcal{L}_{SORS}$ | **Projection Head**<br>FC [1024] + swish<br>FC [512] + swish<br>FC [128] + swish<br><br>$\mathcal{L}_{ssl}$ | **Projection Head**<br>FC [1024] + swish<br>FC [512] + swish<br>FC [128] + swish<br><br>$\mathcal{L}_{ssl}$ | **Classification Head**<br>Normalisation layer<br>FC [128] + swish<br>Dropout<br>FC [4]<br><br>$\mathcal{L}_{SORS}$ |

**Table 1.** Architectures Summary

### 2.2 Whole Slide Classification

A frozen feature extractor (DenseNet121) was applied to extract relevant representations from the tiles. The feature extractor was initialized sequentially with different pre-trained weights. The obtained sets of features were used to train the Attention-MIL model with SORS loss. At each epoch, 400 tiles representations were sampled from each slide to represent the slide in the batch. We used a batch size of 16 and trained the model for 200 epochs, keeping the best weights that maximized the Weighted Accuracy on the validation set. We used the RMSprop optimizer (learning rate: 5e-5, momentum: 0.5) and SORS loss. Dropout layers with rate 0.7 were inserted between the dense layers to limit overfitting, see summary table 2.

| **Layers** | **Type** |
|---|---|
| **Feature Extractor** | DenseNet121 [1024] |
| **Attention-MIL** | **Dimensionality Reduction**<br>FC [128] + tanh<br>Dropout<br>**Tile Scoring**<br>FC [128] + softmax<br>Dropout<br>**Classification**<br>FC [200] + relu<br>Dropout<br>FC [100] + relu<br>Dropout<br>FC [4] |

**Table 2.** Whole Slide Classification - Summary Table

#### 2.3 Data Augmentation

**Self-supervision** See table 3

| Type | Parameters |
|---|---|
| crop and resize | resize to 224x224<br>This augmentation also allows to augment the data with translation<br>Use distorted bounding box crop with parameters:<br>• min_object_covered=0.7<br>• aspect_ratio_range=(3. / 4 , 4. / 3.)<br>• area_range=(0.7, 1.0)<br>• max_attempts=100 |
| random rotation | 0, 90, 180, 270 degrees |
| random flip | up and down<br>left and right |
| color jittering | • brightness = 0.25<br>• contrast = 0.25<br>• saturation = 0.25<br>• hue = 0.1 |
| stain augmentation | according to (Tellez et al. 2018)<br>alpha sampled in [0.975, 1.025]<br>beta sampled in [-0.025, 0.025] |

**Table 3.** Data augmentation parameters for self-supervised model

**Full-supervision** See table 4

| Type | Parameters |
|---|---|
| blur | max sigma: 4 |
| random rotation | max angle 30 degrees |
| random flip | up and down<br>left and right |
| color jittering | • brightness = 0.25<br>• contrast = 0.4<br>• saturation = 0.4<br>• hue = 0.1 |
| translate | max shift: 5 pixels |
| zoom | • max zoom out: 0.75<br>• max zoom in: 1.25 |

**Table 4.** Data Augmentation parameters for fully supervised model

#### 2.4 Feature visualization generation

The linear model was trained for 200 epochs with categorical cross entropy and L1 regularization on the weights (0.1). ADAM optimizer was used with a learning rate of 1e-3. Feature visualizations were generated using open-access re-implementation of Lucid (*https:github.com/pokman/mini_lucid_tf2*, inspired from *https:github.com/tensorflow/lucid* ) and for each feature, optimization was performed for 200 epochs (ADAM optimizer, learning rate of 0.01). The gradient descent steps were made in the decorrelated space and used a frequency decay of 1. Random crops, random rotations and the H&E space constraint were applied on the input for all sets of pre-trained weights except for ImageNet, since it was trained on natural images.

### 3 Additional Results

#### 3.1 $\beta$ optimization

To determine the best $\beta$ parameter we trained the mixed-supervised model for several values of $\beta$ (0, 0.3, 0.7, 0.95) on the 3 tiles splits and used the obtained weights to train the whole slide classification of the 3 respective slides splits. The best Weighted Accuracy was obtained for $\beta = 0.3$. To certify the benefit of our method we also compared the training and validations losses and observed that for $\beta = 0$ (ie. training only the supervised branch) the SSL loss was increasing over epochs, demonstrating that without joint optimization of the two losses, the model was forgetting features learned for the SSL task.

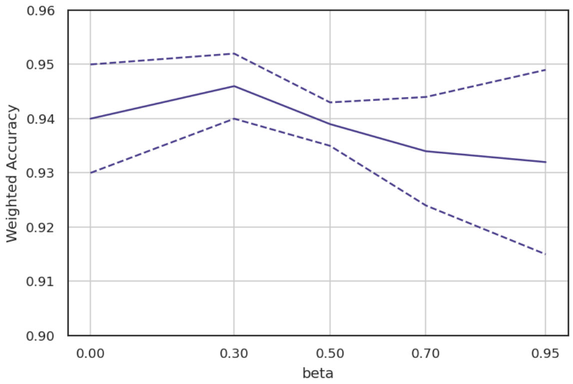


**Fig. 1.** $\beta$ **Parameter Choice** - Weighted Accuracy mean and standard deviation on 3 folds CV for WS classification for different values of $\beta$

#### 3.2 Lasso Training Performances

| Pre-training Policy | Weighted Accuracy |
|---|---|
| ImageNet | 0.865 |
| Supervised | 0.941 |
| SSL | 0.897 |
| Mixed | 0.965 |

**Table 5.** Weighted Accuracy of Lasso training measured on set of test tiles

#### 3.3 Features Visualizations

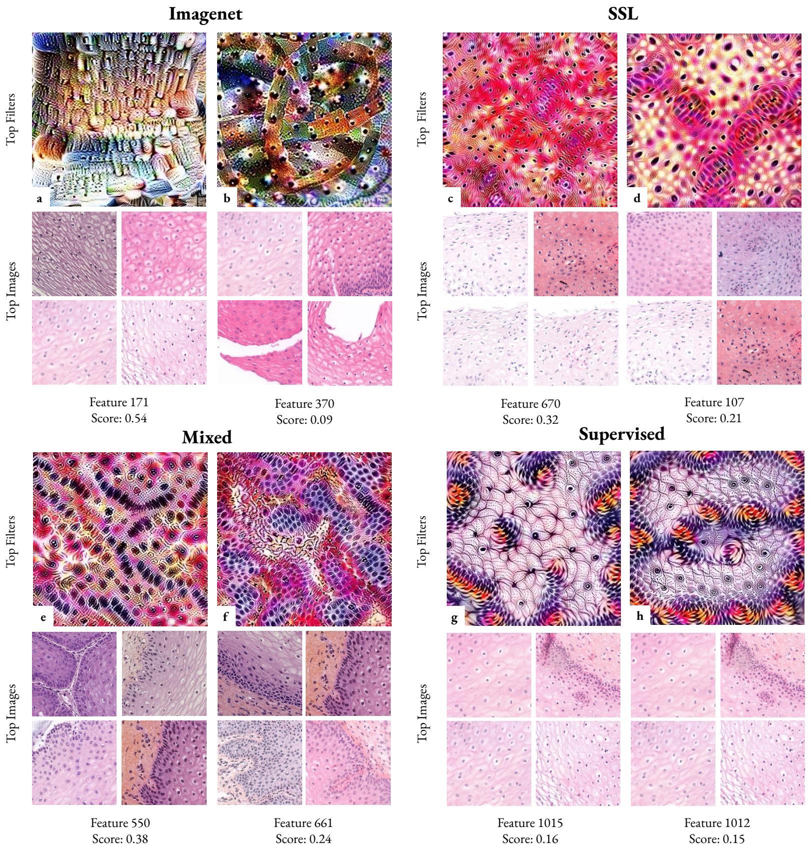


**Fig. 2.** Features visulization for class "Normal" (0)

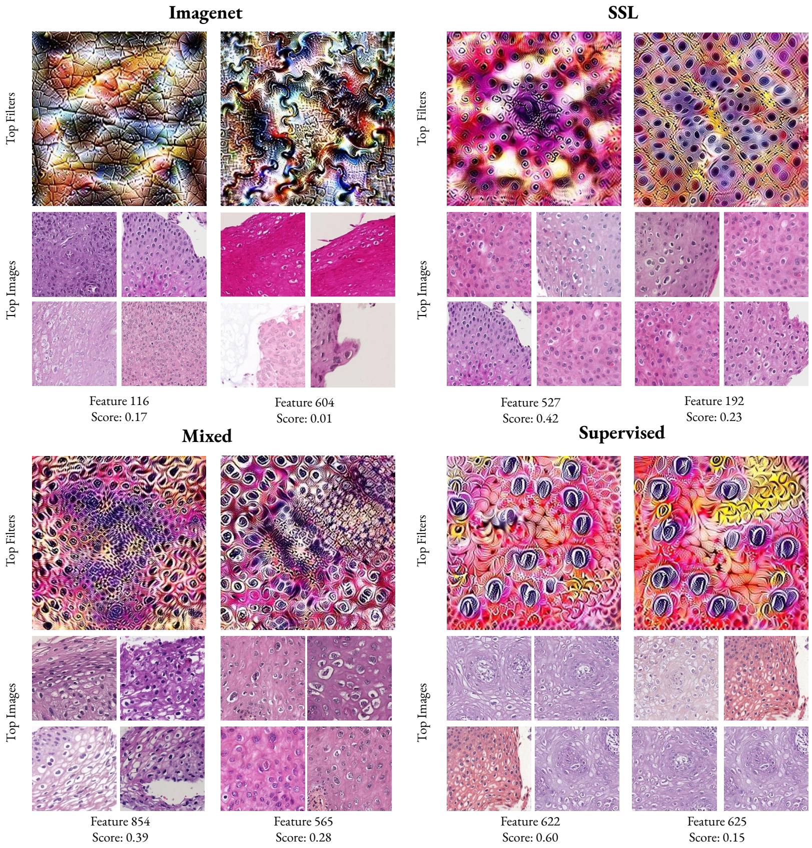


**Fig. 3.** Features visulization for class "Low Grade" (1)

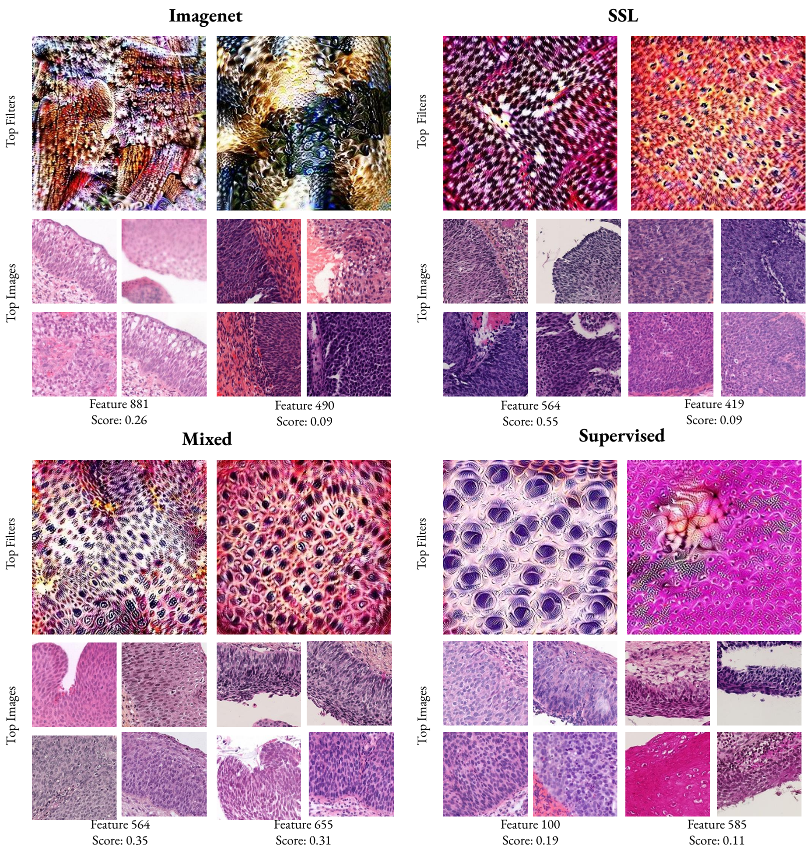


**Fig. 4.** Features visulization for class "High Grade" (2)

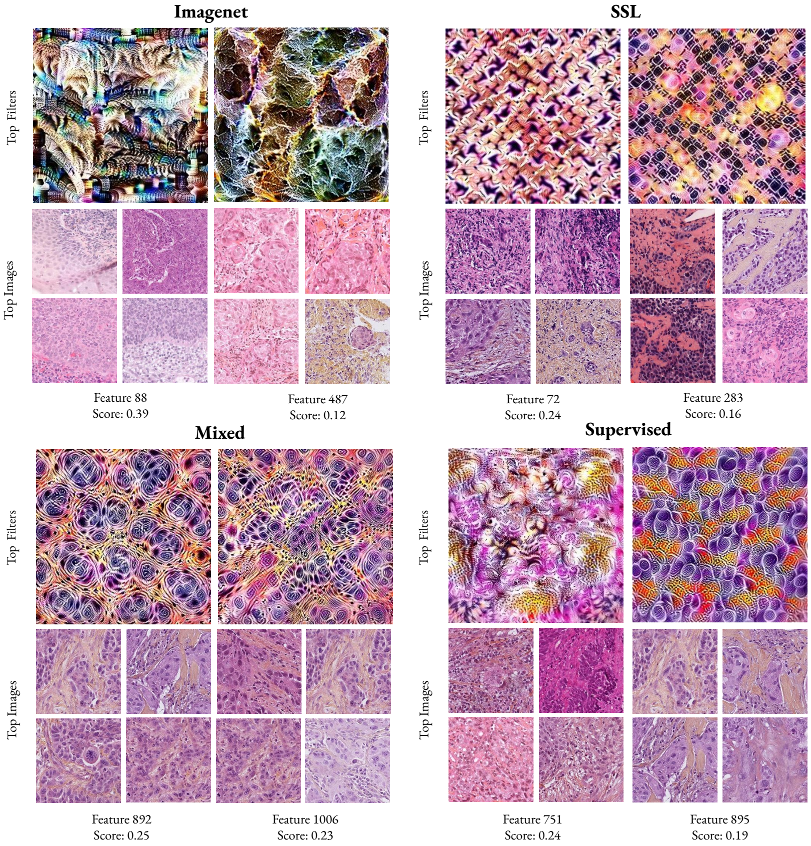


**Fig. 5.** Features visulization for class "Carcinoma" (3)

#### 3.4 Features Redundancy

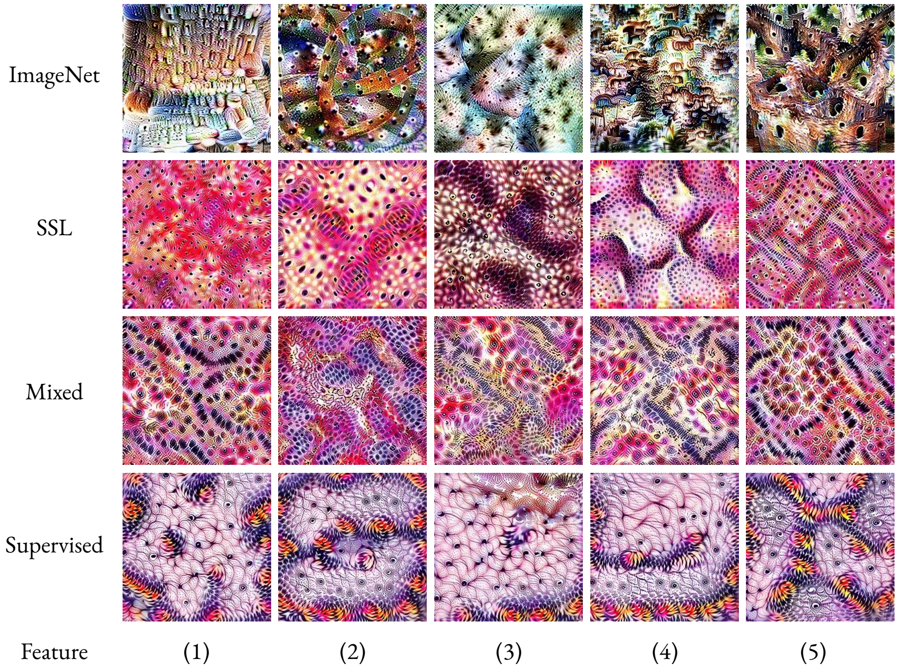


**Fig. 6.** Features Diversity for class "Normal" (0) - Features 1 to 5

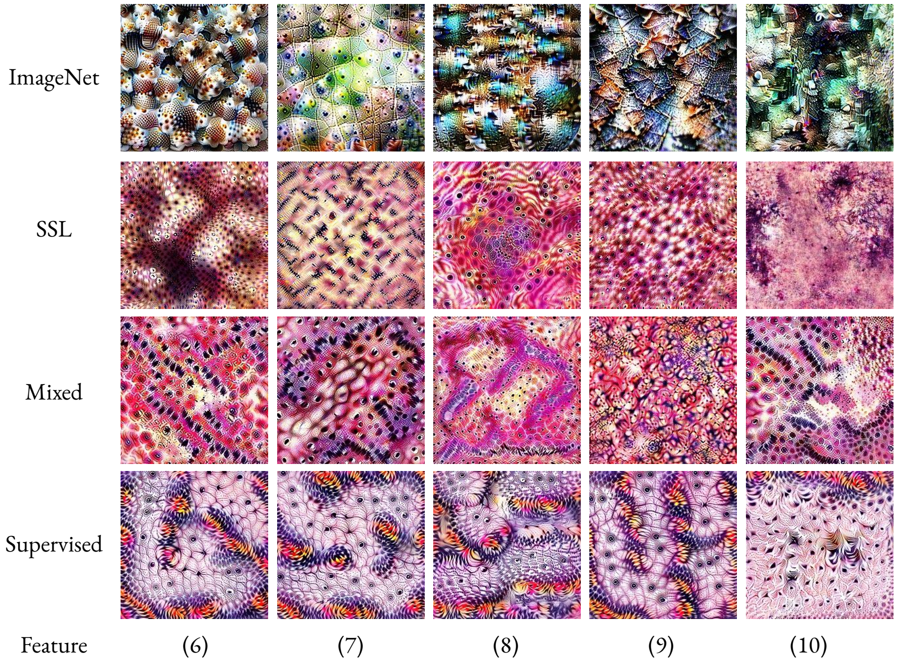


**Fig. 7.** Features Diversity for class "Normal" (0) - Features 5 to 10

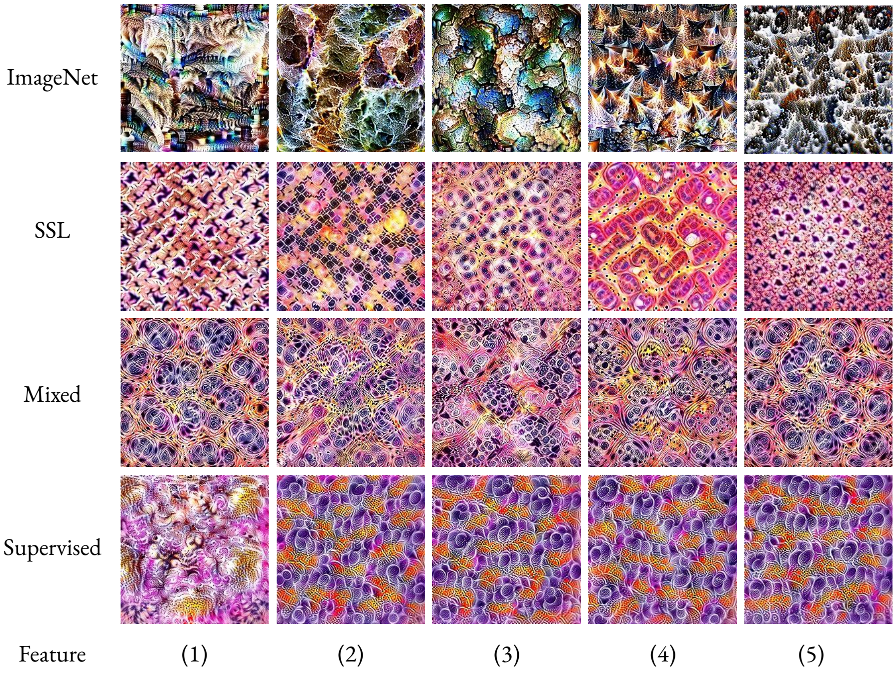


**Fig. 8.** Features Diversity for class "Carcinoma" (3) - Features 1 to 5

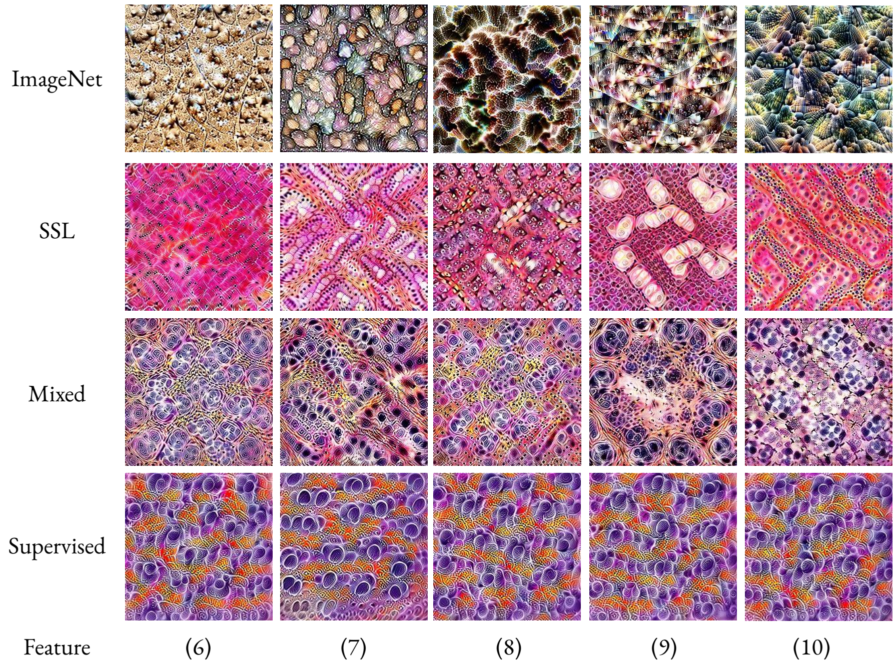


**Fig. 9.** Features Diversity for class "Carcinoma" (3) - Features 5 to 10
